# Endothelial ERG programs neutrophil transcriptome for sustained anti-inflammatory vascular niche

**DOI:** 10.1101/2024.05.02.591799

**Authors:** Vigneshwaran Vellingiri, Vijay Avin Balaji Ragunathrao, Jagdish Chandra Joshi, Md Zahid Akhter, Mumtaz Anwar, Somenath Banerjee, Steven Dudek, Yoshikazu Tsukasaki, Sandra Pinho, Dolly Mehta

## Abstract

Neutrophils (PMNs) reside as a marginated pool within the vasculature, ready for deployment during infection. However, how endothelial cells (ECs) control PMN extravasation and activation to strengthen tissue homeostasis remains ill-defined. Here, we found that the vascular ETS-related gene (ERG) is a generalized mechanism regulating PMN activity in preclinical tissue injury models and human patients. We show that ERG loss in ECs rewired PMN-transcriptome, enriched for genes associated with the CXCR2-CXCR4 signaling. Rewired PMNs compromise mice survival after pneumonia and induced lung vascular inflammatory injury following adoptive transfer into naïve mice, indicating their longevity and inflammatory activity memory. Mechanistically, EC-ERG restricted PMN extravasation and activation by upregulating the deubiquitinase A20 and downregulating the NFκB-IL8 cascade. Rescuing A20 in *EC-Erg^-/-^* endothelium or suppressing PMN-CXCR2 signaling rescued EC control of PMN activation. Findings deepen our understanding of EC control of PMN-mediated inflammation, offering potential avenues for targeting various inflammatory diseases.

**Highlights:**

- ERG regulates trans-endothelial neutrophil (PMN) extravasation, retention, and activation
- Loss of endothelial (EC) ERG rewires PMN-transcriptome
- Adopted transfer of rewired PMNs causes inflammation in a naïve mouse
- ERG transcribes *A20* and suppresses CXCR2 function to inactivate PMNs

**In brief/blurb:** The authors investigated how vascular endothelial cells (EC) control polymorphonuclear neutrophil (PMN) extravasation, retention, and activation to strengthen tissue homeostasis. They showed that EC-ERG controls PMN transcriptome into an anti-adhesive and anti-inflammatory lineage by synthesizing *A20* and suppressing PMNs-CXCR2 signaling, defining EC-ERG as a target for preventing neutrophilic inflammatory injury.

## Introduction

Polymorphonuclear neutrophils (PMNs) are the most abundant type of white blood cells and the first responders to be recruited from circulation to the tissues following environmental and blood-borne infections [1, 2]. These cells eradicate infections by generating reactive oxygen species (ROS), cytotoxic granules, extracellular trap formation, and phagocytosis [3-6]. PMNs are short-lived cells but can accumulate at affected sites in adequate numbers or return from the tissues to the bone marrow by reverse migration [4, 7, 8]. Studies show that other than the bone marrow, spleen, and liver, the lung microvasculature is a crucial reservoir of PMNs, defined as the lung-marginated neutrophil pool [9-11]. The lung-marginated PMN pool can replenish the circulating population or defend against pathogens when needed [4, 10]. PMNs also exhibit functional and phenotypic diversity based on the tissue niche. For example, lung neutrophils are enriched for the G protein-coupled receptor (GPCR) C-X-C chemokine receptor type 4 (CXCR4), which allows neutrophil retention via CXCL12 binding before trafficking back to the bone marrow, spleen, and liver where they are cleared [11-13].

The large surface area of the lung environment is constantly exposed to inhaled pathogens and other environmental stimuli. An abundant PMN resident population in the immediate vicinity ensures proper immune function. However, it also underscores the importance of tightly regulating neutrophil transmigration across the microvasculature to safeguard the underlying tissue from an excessive innate immune response [2, 14]. Indeed, hyperpermeable lung microvasculature with neutrophilic inflammation is consequential to lethal acute lung injury (ALI) and acute respiratory distress syndrome (ARDS) [15-17] due to a lack of therapeutic options [18]. Understanding vascular mechanisms that control PMN extravasation and activation in the lung circulation is critical to improving treatment options for ALI/ARDS and other chronic inflammatory diseases.

The ETS-related gene (ERG) transcription factor can regulate multiple regulatory elements as a super-enhancer [19-21]. The crucial role of ERG in the vasculature was revealed in genetic studies showing that EC deletion of ERG leads to compromised vascular development, induced vascular leak, and increased mice susceptibility to developing fibrosis during aging [22-27]. ERG was initially reported to bind the interleukin-8 (IL8) promoter (MIP2α in mice)[28]. IL8 is a crucial proinflammatory cytokine linked with neutrophil recruitment to tissue and immune disorders, including ALI/ARDS and sepsis [29, 30]. Whether ERG is required for suppressing neutrophil infiltration and activation to strengthen vascular homeostasis remains ill-defined. Here, using mice lacking ERG specifically in the endothelium, we show that ERG is required to maintain PMN transcriptome that limits PMN accumulation, activation, and lung inflammation. Mechanistically, we show that ERG’s function involves synthesizing multipurpose deubiquitinase A20 and suppressing the PMN-CXCR2 signaling.

## Results

### Endothelial ERG suppresses neutrophil infiltration and activation to maintain tissue homeostasis at a steady state

We first evaluated ERG expression across distinct tissues. We found that at steady state ERG protein, as well as mRNA expression levels, were markedly higher in the lung than in the kidney or brain (**Suppl Fig. 1A)**. Moreover, the publicly available data set indicated that ERG was more highly expressed in ECs than other pulmonary cells (**Suppl Fig. 1B**). Thus, we focused our studies on the lung as a model to study the role of ERG in regulating immune cell functions because the pulmonary microvasculature acts as a crucial reservoir of PMNs, and hyperactivation of neutrophils causes lethal inflammatory lung injury [2, 4, 14]. To this end, we bred ERG floxed mice (*Erg^fl/fl^*) with the tamoxifen-inducible *Cdh5^Cre-ERT2^* line to generate *Erg-Cdh5^Cre-ERT2^* mice and induced conditional deletion of ERG in EC, hereafter referred to as *iEC-Erg^-/-^*(**Fig. 1A**). As expected, 11 days after tamoxifen treatment, Cre recombinase activity markedly decreased *Erg* mRNA levels in sorted ECs without altering the expression of the closely related ETS transcription factor, *Fli1* (**Suppl Fig. 1 C**). Immunoblot and confocal imaging confirmed ERG deletion in pulmonary vessels (**Suppl Fig. 1 D-E).** In agreement with previous studies [31], our results showed that EC-ERG null lungs were also inflamed as the mRNA expression levels of inflammatory cytokines (*IL6*, *Il1β*, *Mip2α,* and *Tnfa*) (**Suppl Fig. 1F)** and myeloperoxidase (MPO) activity (**Suppl Fig. 1G)** were also increased. Notably, ECs lacking ERG at homeostasis also showed increased *Mip2α* and *Tnfα* transcripts than in controls (**Fig. 1B)**.

**Figure 1:**
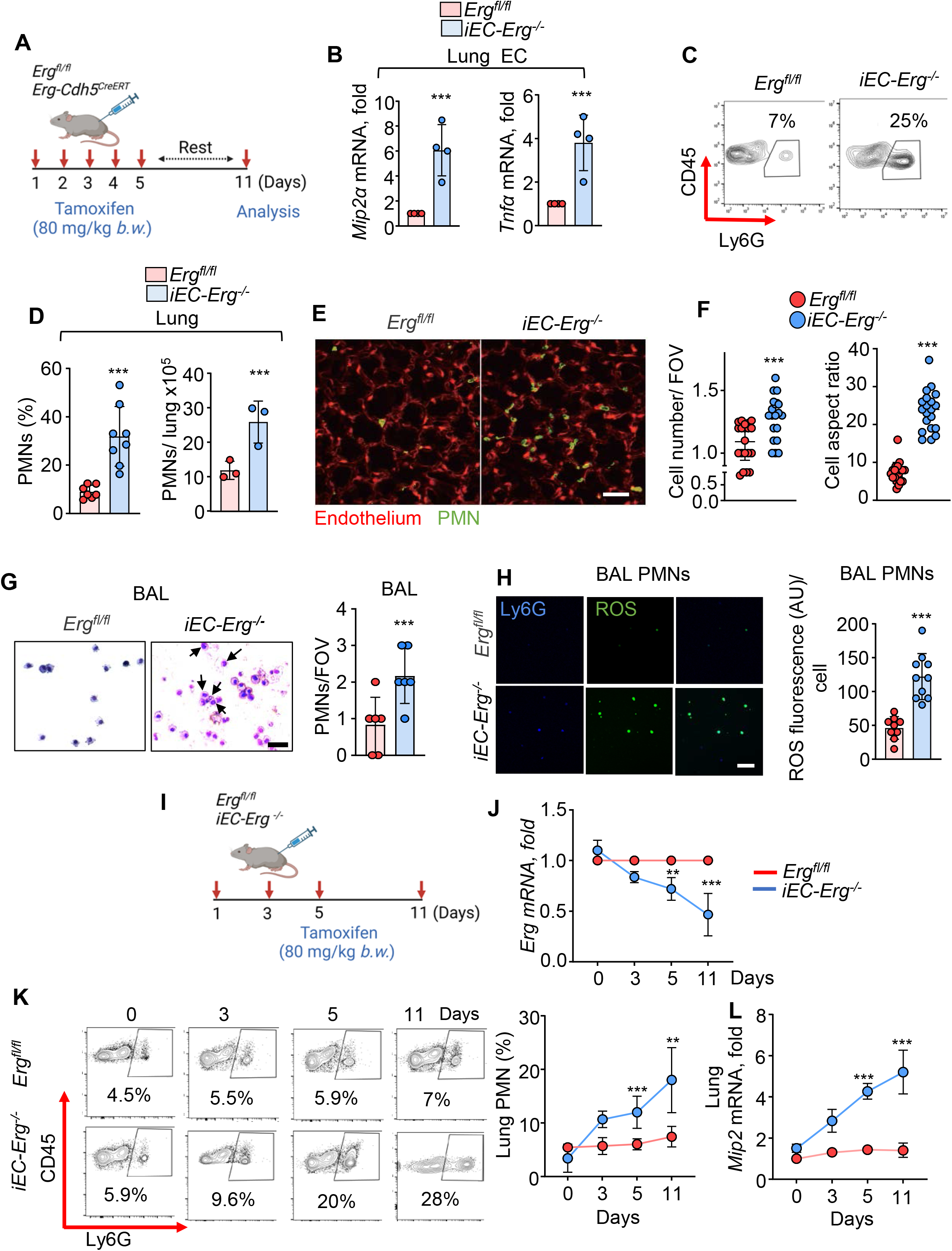
Endothelial-ERG loss induces neutrophil infiltration and activation. **(A)** Schematics of ERG deletion in EC and assessment of PMNs functions. **(B)** *Mip2αα* and *TNFα* mRNA in EC sorted from the lungs of indicated mice. GAPDH was used as an internal control. **(C-D)** Flow cytometric analysis of PMNs in indicated lungs. A representative FACS plot is shown in **C**, while **D** shows PMN percentage (left) and absolute count (right) (n=6). (**E and F)** Intravital two-photon analysis of PMNs (green) and EC (purple) in the indicated lungs following *i.v.* injection of BV421-Ly6G (for PMNs) and Setau647 CD31 (for ECs) antibodies. The lungs were imaged after 30 min. Scale bar = 50 μm. A representative image is shown in **E**, while **F** shows an analysis of PMN cell number and aspect ratio (Field of view, FOV, is 298 x 298 mm) (n=6). Experiments were performed two times independently. (**G**) Hematoxylin and eosin-stained BAL from indicated mice showing airspace infiltrated PMNs. The left is a representative image, while the right shows the quantitation of PMNs/FOV. Scale bar = 100 μm. (**H**) ROS levels in BAL PMNs from indicated mice after staining with Ly6G antibody and H2-DCFDA. A representative image is shown on the left, while the right shows quantitation. **(I**) Schematics of the time course of PMN accumulation in lungs upon ERG deletion in EC. (**J**) Lung ERG mRNA in indicated mice taking GAPDH as control (n=6). (**K**) A representative FACS plot (left) and PMNs analysis (right) (n=3). (**L**) Lung *Mip2α* mRNA taking GAPDH as controls (n=3). Data represented as ±SEM. Unpaired Students’ t-tests (for figures B, D, F, G, and H). One-way ANOVA followed by Tukey’s multiple comparison tests (J, K, and L). P*<0.05, **<0.01, ***<0.001, ****<0.0001.

To investigate the effect of EC-ERG deletion on lung immune cells, we performed flow cytometry analyses of control *Erg^fl/fl^* and *iEC-Erg^-/-^* lungs first using CD31 (a canonical EC marker) and CD45 (a pan-hematopoietic marker) antibodies (**Suppl Fig. 2A**). Compared to *Erg^fl/fl^* lungs, *iEC-Erg^-/-^* lungs showed reduced EC number (%) (CD45^-^CD31^+^) while CD45^+^ hematopoietic cells were increased (**Suppl Fig. 2A-C**). Using cell-surface markers of myeloid (IMs, CD45+CD64+CD11b+, AMs CD45+CD64+SiglecF+, PMNs, CD31-CD45+Ly6G+), platelets (CD45+CD62P+), and lymphoid cells (CD45+CD3+), we found a ∼7-fold increase in PMN accumulation in the EC-ERG null lungs but not of macrophages, platelets or lymphocytes (**Fig. 1C-D and Suppl Fig. 2D)**. Accordingly, we observed an increase in the frequency of PMNs in circulation, but not in the bone marrow of *iEC-Erg^-/-^* mice compared to control (**Suppl Fig. 2E**).

The enhanced PMN cell number in ERG null lungs and circulation could be due to increased cell proliferation. However, Ki67 and Annexin staining of lung PMNs did not reveal any differences **(Supp Fig. 3A).**

To determine whether EC-ERG null lung neutrophils acquire functions distinct from control neutrophils, such as migratory, extravascular, and phagocytic [1, 4], we performed intravital two-photon imaging of lungs. We generated ERG-Cdh5^CreERT2^; R26R^t*dTomato*^ mice to allow “Red” labeling of ECs following tamoxifen-induced ERG deletion. These mice received Ly6G antibodies for PMN labeling before imaging (**Fig. 1E**). In agreement with our flow cytometry analyses, we observed a markedly increased neutrophil cell number in EC-ERG null lungs (**Fig. 1E-F**). As expected, PMNs rushed rapidly around the vasculature in control lungs, but in contrast, PMNs spread out more and adhered to the vasculature in EC-ERG null lungs **(Fig. 1F)**. Consistently, bronchoalveolar lavage (BAL) showed a markedly increased PMN cell number in the airspace of EC-ERG null lungs but not in control mice (**Fig. 1G**).

Neutrophils can phagocytize and kill pathogens by generating ROS [5, 6]. EC-ERG PMNs produced markedly higher ROS levels than controls (**Fig. 1H**). Interestingly, while BAL neutrophils from EC-ERG null lungs showed poly-segmented nuclei, they also presented donut-shaped nuclei, which may augment inflammation seen in EC-ERG lungs (**Suppl Fig. 3B**). To compare the phagocytic activity of PMNs we used *E. coli* bioparticles and then we lavaged PMNs from EC-ERG null lungs. However, due to limited PMN infiltration in normal lungs, we used PMNs isolated from the lungs of *Erg^fl/fl^* mice after the LPS challenge to compare the phagocytic activity of PMNs from EC-ERG null lungs. EC-ERG null PMNs phagocytized bacteria as efficiently as LPS-exposed control neutrophils (**Suppl Fig. 3C**), confirming their hyper-activated state.

These results prompted us to the relationship between ERG deletion in EC with the increase in neutrophil cell number in the lungs (**Fig. 1I**). Tamoxifen-induced ERG decreased in EC by 80% on day 4^th^, which further reduced to almost 95% on day 5^th^ and sustained at this level at 11 days (**Fig. 1J**). Interestingly, 80% deletion of EC-ERG increased lung PMNs by 2-fold on day 3rd, whereas 95% of ERG deletion increased PMNs by 3-fold on days 5^th^ and 11^th^ (**Fig. 1K**). These differences translated into increased lung inflammation as reflected by a day-wise increase in *Mip2α* expression in EC-ERG null lungs (**Fig. 1L)**.

Next, we assessed the impact of EC-ERG deletion-associated neutrophilia on lung homeostasis. Consistent with previous findings [31], mice lacking ERG in EC developed fulminant lung edema (**Suppl Fig. 3D**) and lung injury as revealed by marked septal thickening and leukocyte infiltration (**Suppl Fig. 3E**). To corroborate the clinical relevance of the above findings, we also analyzed ERG expression in human lung vessels of patients who died of ARDS compared to patients dying of other non-lung-related causes. We also observed a significant reduction in ERG expression levels and PMN activation, as reflected by increased MPO activity, in the lungs of patients with ARDS (**Suppl Fig. 3F-G**). Importantly, human data was consistent with our murine studies. These results, therefore, identify vascular ERG as the instructor of the anti-inflammatory vascular niche as its loss subverted EC to adapt inflammatory lineage specifically targeted for promoting PMN extravasation, accumulation, and activation, leading to neutrophilic vascular injury.

### Neutrophils transcriptome is rewired into an inflammatory program in EC-ERG-deleted lungs

PMN transcriptome is highly malleable during circadian variation, cancer metastasis, cardiovascular diseases, and autoimmune diseases [13, 32-35]. To address if inflammatory EC niche rewired PMN transcriptome, we performed panRNAseq of PMNs sorted from *iEC-Erg^-/-^* and *Erg^fl/fl^* control lungs (**Figure 2A**). Heatmap and volcano plot from RNA sequencing showed that out of 3944 genes, 1590 genes were differentially expressed in the neutrophils from *iEC-Erg^-/-^* lungs than control lungs (**Fig. 2B and Suppl Fig. 4A**). Ingenuity Pathway analysis of the top 100 transcripts showed that *iEC-Erg^-/-^* PMNs are enriched with genes inducing an inflammatory response, negative regulation of apoptosis, migration, and chemotaxis, corroborating the above functional findings **(Fig. 2C and Suppl Fig. 4B)**. While screening for the top 10 upregulated genes in EC-ERG null PMNs, we found increased expression of genes associated with pro-inflammation and delayed apoptosis (**Figure 2D**). We next validated by qPCR the increased expression of some of these critical genes, such as *Il1β*, *Tnfα, Cxcr2, Cxcr4, and Marcksl1* in PMNs sorted from EC-ERG null lungs (**Figure 2E)**. We also found increased cell-surface expression of CXCR2 and CXCR4, known to be critical for PMN retention in the lung [4](**Fig. 2F and Suppl Fig. 4B**). Consistently, EC-ERG null showed CXCR2 and CXCR4 expressing populations in the blood as analyzed by FACS **(Suppl Fig. 4 C).**

**Figure 2:**
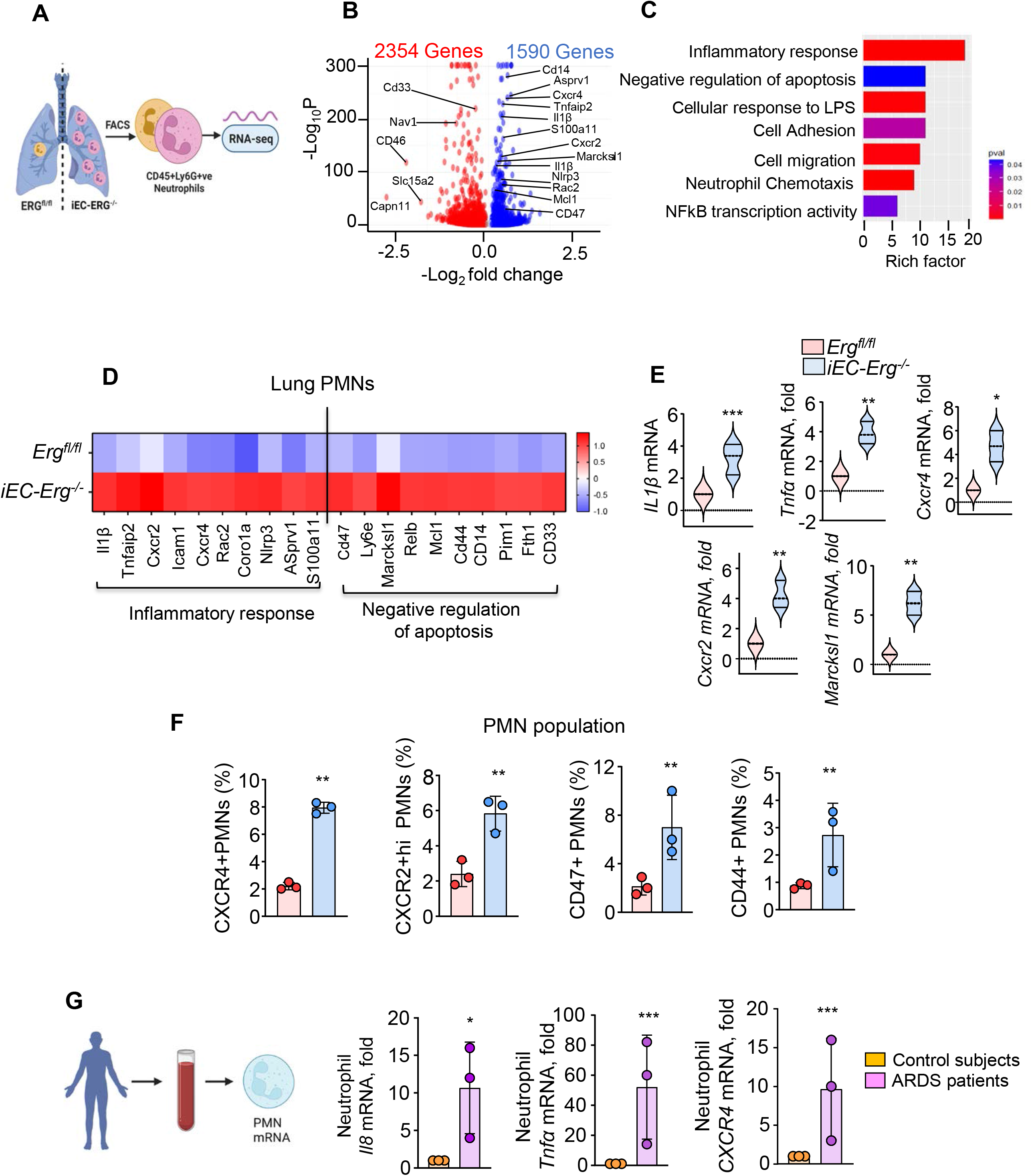
Neutrophil transcriptome is shaped by EC-ERG. (**A**) Schematics of lung PMNs sorting for RNA sequencing. (**B-D**) volcano plot (**B**) out of which top 100 genes were selected for pathway enrichment analysis (**C**) and **(D)** heat map. **(E)** Validation of PMNs mRNA expression using GAPDH as an internal control (n=3). (**F**) Cell surface expression of CXCR4, CXCR2, CD47, and CD44 in the indicated PMNs using flow cytometry. (n=3). (**G)** mRNA expression of indicated genes in human peripheral blood neutrophils from normal subjects and ARDS patients (n=3). Data represented as ±SEM. Unpaired Students t-test (for Figs. E, F, and G), where P *<0.05, **<0.01, ***<0.001, ****<0.0001.

CD47 expression in PMNs is associated with a delay in neutrophil apoptosis and phagocytic clearance[36]. CD44 is a crucial cell-surface receptor on PMNs that regulates leukocyte extravasation into an inflammatory site upon binding with hyaluronic acid (HA) [37]. Notably, FACS analysis showed that the frequency of CD44+ and CD47+ PMNs population was increased three-fold in EC-ERG lungs (**Fig. 2F and Suppl Fig. 4D**).

To evaluate whether increased expression of *Il8* (mouse Mip2α), *Cxcr4, and Cxcr2* expression is correlated clinically, we measured the expression of these genes in the PMNs sorted from the blood of ARDS patients in the ICU versus control subjects. ARDS patients exhibited augmented mRNA expression of *Il8, Tnfα,* and *Cxcr4* in blood PMNs compared to the controls (**Fig. 2G)**. These results, therefore, identify prominent changes in the PMN transcriptome associated with their extravasation, residence, and lung inflammation, which are also evident in the blood of ARDS patients.

### Rewired neutrophils sorted from EC-ERG lungs retain inflammatory memory in a naïve recipient

Evidence suggests that *Cxcr4* is highly expressed in aged and inflammatory neutrophils and that increased *Cxcr2/Cxcr4* expression promotes cell migration and lung retention [4]. Therefore, we adoptively transferred EC-ERG PMNs alongside control PMNs (1.0 x10^6^ *i.v.*) into LyZ-M-GFP reporter mice in which all myeloid cells are fluorescently labeled **(Figure 3A)** and determined lung PMN frequencies, endothelial injury, and inflammatory gene expression four hours post-transplantation. Notably, the frequency of donor EC-ERG null PMNs (GFP^-^) was 2-fold higher than control donor PMNs in the lungs of naïve recipient mice **(Figure 3B).** The frequency of pulmonary GFP^+^ recipient PMNs was unaffected **(Figure 3B)**. Notably, EC-ERG null transplanted neutrophils led to inflammatory lung injury in naïve recipient mice, as indicated by increased lung edema and inflammatory cytokines compared to control (**Fig. 3C-D**). These findings suggest that PMNs from EC-ERG null lungs were wired for the longevity and memory of their inflammatory activity.

**Figure 3:**
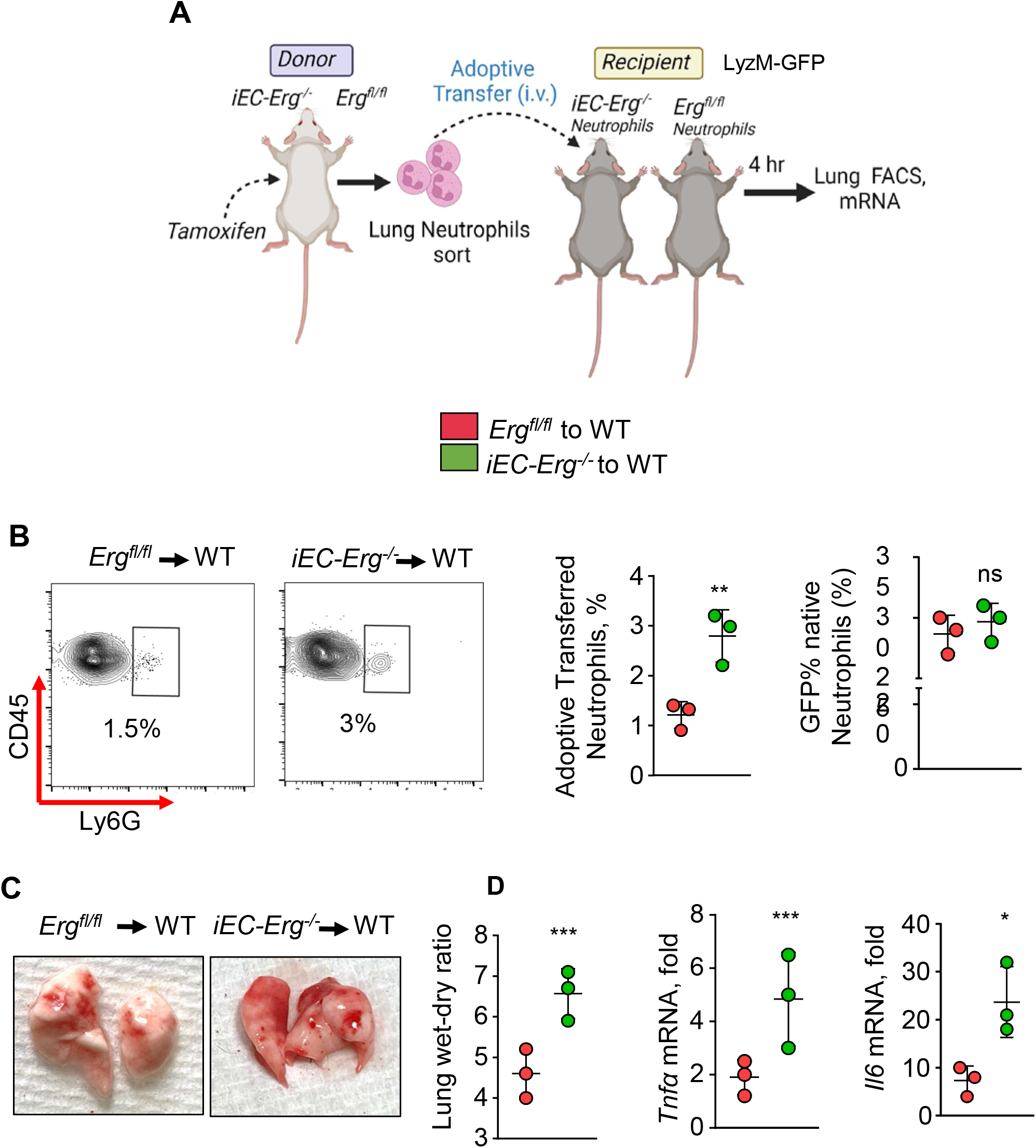
Adoptively transferred neutrophils from EC-ERG null mice induces lung injury. **(A)** Schematic of PMNs adoptive transfer. **(B)** FACS analysis of adoptively transferred neutrophils in the lungs (n=3). The left shows the FACS plot, while the right shows quantitation. n=3. (**C)** Lung wet-dry ratio in the recipient mice. The left shows the representative image, while the right shows the quantitation. **(D)** mRNA expression of indicated genes in the recipient’s lungs using GAPDH as an internal control (n=3). Data represented as ±SEM. Unpaired Students t-tests (for B, C, and D). P*<0.05, **<0.01, ***<0.001, ****<0.0001.

### EC-ERG controls neutrophilic lung injury by transcribing A20

NFκB is a crucial transcription factor inducing inflammatory signaling downstream of diverse pathogens in EC [12]. Next, we tested the hypothesis that ERG controls PMN’s activation by suppressing NFκB activity [38]. We found a 5-fold increase in NFκB phosphorylation, a measure of NFκB activity [39] in EC-ERG null lungs compared to control lungs (**Figure 4A)**. Also, EC-ERG null lungs showed markedly enhanced NFκB and IKKα protein expression and VCAM1 and IL6 pro-inflammatory molecules mRNA expression (**Figure 4A-B**). *Erg* depletion in human ECs similarly induced NFκB signaling (**Figure 4C-D and Suppl 5A**), suggesting that ERG suppression of NFκB is a conserved mechanism. These findings demonstrate that ERG represses basal NFκB activity to maintain an anti-inflammatory vascular niche.

**Figure 4:**
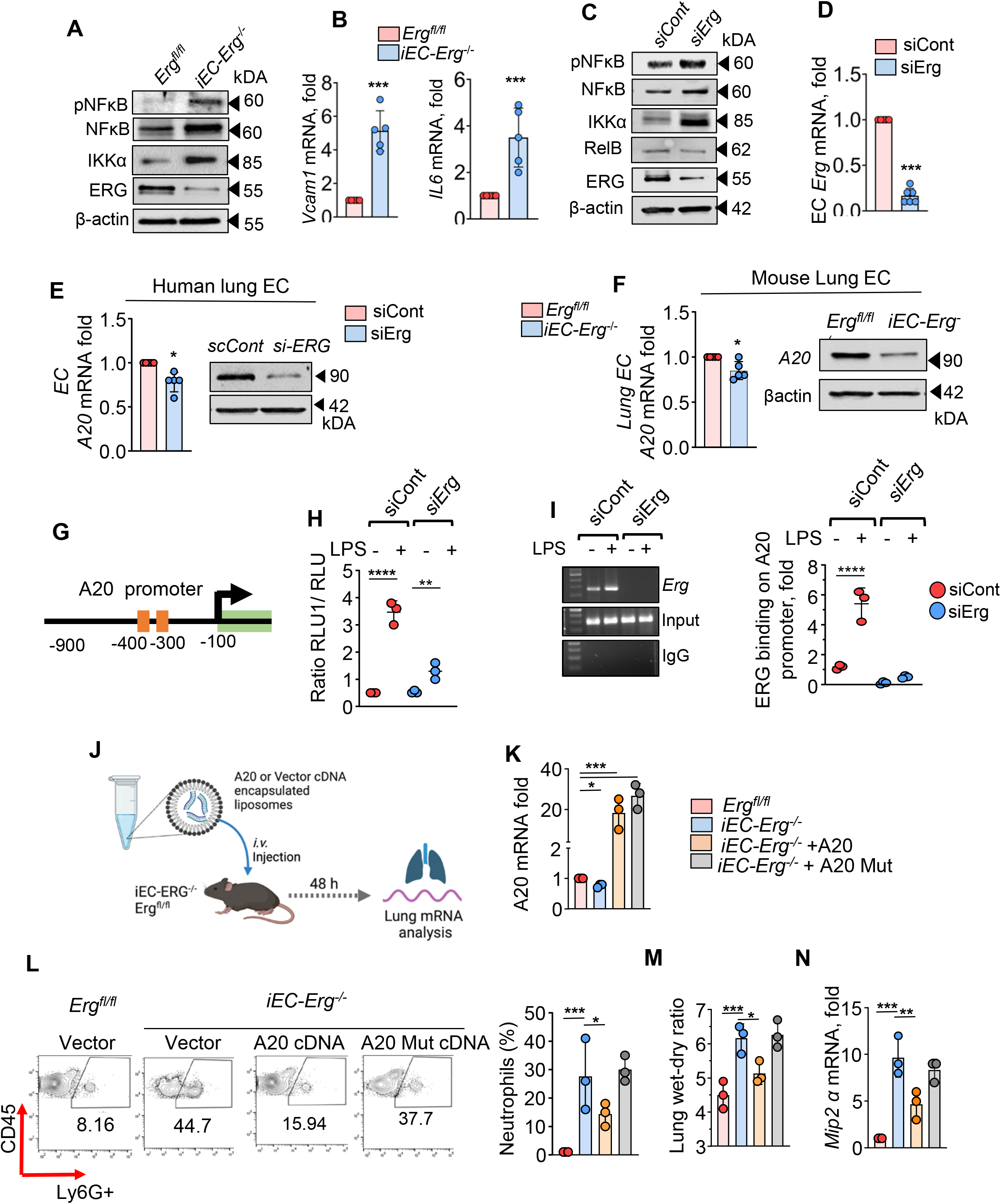
ERG suppresses PMN infiltration and activation by synthesizing A20. **(A)** Immunoblot of indicated proteins from lung lysates. Beta-actin was used as a loading control. A representative blot is shown from experiments that were repeated three times. **(B)** mRNA expression of indicated genes taking GAPDH as an internal control (n=6). **(C)** Immunoblot of indicated proteins in ERG-depleted ECs. Actin was used as a loading control. A representative blot is shown from the experiment. **(D).** ERG mRNA expression in indicated ECs. GAPDH was used as an internal control for mRNA. **(E-F)** A20 mRNA (left) and protein (right) expression in human lung ECs (**E**) and mouse lung ECs (**F**). GAPDH was used as an internal control for mRNA (n=5), while actin was used as a loading control for protein. A representative immunoblot is shown from experiments that were repeated three times. **(G)** In silico analysis of A20 promoter showing two ERG binding sites. **(H)** A20 promoter activity. **(I)** *Chip* of A20 with ERG. The left shows a representative blot, while the right shows the quantitation of PCR’s product. **(J)** Schematics of liposomal gene delivery. **(K)** Lung A20 mRNA expression using GAPDH as an internal control. n=3 lungs/group. **(L)**. PMNs analysis. A FACS plot is shown along with the quantitation. n=3. (**M**). Lung edema determined by quantifying the lung wet-dry ratio. (**N**) Lung *Mip2α* mRNA taking GAPDH as the loading control. Data represented as ±SEM. Unpaired Students’ t-tests (for Figs. B, D, E, and F) and One-Way ANOVA followed by Tukey’s multiple comparison tests (for Figs. H, I, K, L, M, and N). P*<0.05, **<0.01, ***<0.001, ****<0.0001.

We then sought to identify the molecular basis of ERG suppression of the NFκB-mediated inflammation. The deubiquitinase interface with and modify ubiquitylated protein substrates in multiple ways to curb persistent NFκB activation and prevent harm to the host [40, 41]. We screened for deubiquitinates, which are known to be expressed in the lungs and to regulate NFkB signaling (Cyld, USP9X, USP18, and A20 (also known as TNFAIP3) [41]. We found that ERG depletion in human lung ECs reduced A20 mRNA levels by ∼30% without altering the expression of other deubiquitinases (**Figure 4E Suppl Fig. 5B**). Depletion of ERG reduced A20 protein levels by 80% (**Fig. 4E**). Next, we sorted EC from control and EC-ERG deleted lungs and similarly found significantly reduced A20 mRNA expression and protein levels (**Figure 4F**).

LPS increases A20 expression, and loss of A20 in EC induces vascular inflammatory injury [42]. *In silico* analysis showed that ERG has two binding sites within 1 kb of the A20 promoter (**Figure 4G**). We, therefore, transduced the A20 luciferase promoter in control and ERG-depleted EC and found that LPS failed to increase A20 promoter activity in ERG-depleted EC **(Figure 4H)**. Chromatin immunoprecipitation (*ChIP*) analysis with quantitative real-time PCR showed that LPS increased ERG binding to A20 promoter in control cells but not ERG-depleted EC **(Figure 4I).**

To assess whether A20 deubiquitinase function was required in reversing the inflammatory phenotype in ERG-depleted human lung EC, we transduced WT-A20 or A20-cDNA lacking deubiquitinase activity **(Supplementary** Fig 5C). We found that rescuing A20 expression reversed *Il8, Tnfa,* and *Vcam1* pro-inflammatory molecules mRNA to the levels seen in control cells, but this effect was not seen in the EC-transducing A20 mutant (**Supplementary** Fig 5D). We, therefore, assessed whether restoring A20 expression in EC of EC-ERG null mice would reduce neutrophilic inflammation and resolve vascular injury in i*EC-Erg^-/-^* mice. We complexed WT-A20 or A20-dub mutant cDNA driven by the *Cdh5* (VE-cadherin) promoter in liposomes [40, 43] to express *A20* only in EC of i*EC-Erg^-/-^* mice (**Fig. 4J**) and quantified neutrophil cell number, inflammatory cytokines expression and lung edema (**Fig. 4K-N**). Transducing WT-A20 but not the mutant in EC of EC-ERG-null lungs suppressed neutrophil accumulation **(Fig. 4L)**. Moreover, A20, but not the mutant, reduced inflammatory cytokine expression and promoted the resolution of lung edema **(Fig. 4M and N)**. Thus, these studies defined A20 as a novel ERG target via which ERG ensures tissue homeostasis by suppressing neutrophil recruitment and activation.

### Inhibiting PMNs CXCR2 activity subverts inflammatory injury in EC-ERG null mice

IL8 (MIP2α) induces PMN egress from bone marrow and activation through binding its receptor *Cxcr2* [8] (**Suppl Figure 6A**). Reparixin is an allosteric inhibitor of *Cxcr2* [44]. To assess if CXCR2 regulated PMN activity and function in EC-ERG null mice, we depleted ERG in vitro using siRNA and co-cultured with PMNs derived from HL-60 cells [37] without or with Reparixin treatment (**Suppl Fig 6A**). We found ∼5-fold higher PMN attachment to ERG-depleted EC than in control ECs (**Suppl Fig 5B)**. These PMNs showed augmented NFκB signaling as indicated by increased expression and phosphorylation of NFkB, *Tnfa*, and *Cxcr2* mRNA expression (**Suppl Fig. 5C-E**). Blocking *Cxcr2* activity in PMNs suppressed these changes in PMNs (**Suppl Figure 5E**).

Having identified that Reparixin blocked NFκB signaling and cytokine expression in PMNs co-cultured with ERG-depleted ECs, we next injected Reparixin along with tamoxifen in *Erg^—^cdh5^Cre-ERT2^*mice, following which PMN cell numbers and lung inflammation were determined (**Fig. 5A**). Notably, Reparixin restored neutrophil cell numbers in EC-ERG null lungs to the control levels (from around 28% to 14% respectively) (**Figure 5B**). Histopathology examination of lungs using H&E and MPO immunostaining showed a reversal of neutrophilic injury and lung edema in EC-ERG null mice upon Reparixin treatment (**Figure 5C-E).** These observations indicate that CXCR2 signaling in PMNs is a crucial mechanism underlying lung damage in *iEC-Erg^-/-^* mice.

**Figure 5:**
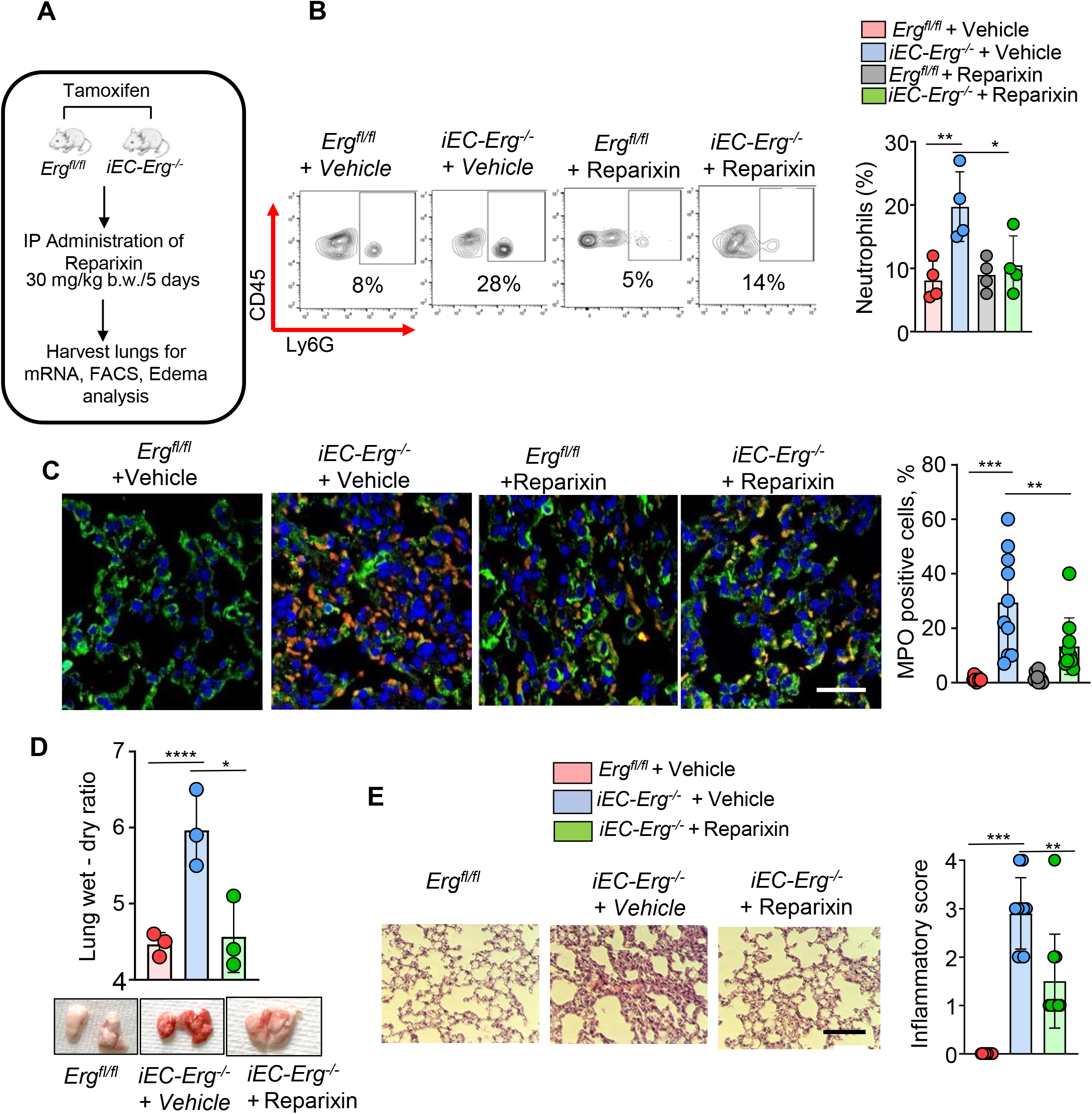
CXCR2 antagonism resolved lung injury in EC-ERG null mice. **(A)** Schematic of Reparixin administration and assessment of PMN functions and lung injury. **(B)** PMNs analysis in the lungs of indicated mice after without or with treatment with Reparixin. The left shows the FACS plot, while the right shows quantitation (n=4). (**C**) Myeloperoxidase (MPO) immunostaining. The left shows a representative image, while the right shows quantitation. Scale bar 20 μm (n=6). (**D)** Lung wet-dry ratio. *iEC-Erg^-/-^* lungs. The top shows the quantitation, while the bottom shows hemorrhagic lungs (n=3). **(E)** Lung H&E staining (left) and the corresponding inflammatory score are shown on the right. (n=6). Data represented as ±SEM. One-way ANOVA, followed by Tukey’s multiple comparison tests (for Fig. B, C, and E). P*<0.05, **<0.01, ***<0.001, ****<0.0001.

### Bacterial pneumonia suppresses EC-ERG expression during injury

Bacterial infection by Gram-positive--bacteria, such as *Pseudomonas aeruginosa* (*PA*), in hospitalized patients, causes ALI, compromising their survival [45, 46]. We, therefore, used a well-characterized mouse model of lung injury where *PA*-induces severe ALI within 12-48 h, which is then resolved in the next 4-5 days (*PA, 1x10^4^ CFU i.t) [47]* **(Figure 6A)** to determine the time course of ERG expression during injury and repair. *PA* infection reduced ERG expression levels by ∼50% at the mRNA and protein levels 12 hours post-infection in the lung (Fig 6B-C) and by ∼80% 24 hours after **(Fig 6B-C)**. Interestingly, ERG expression reversed towards basal levels within 48 to 120 hours (**Fig. 6B-C**). However, we did not observe any change in the expression of other ETS transcription factors, such as Fli-1 **(Fig 6D)**. In agreement with our previous observations, reduced ERG expression in pulmonary vessels was associated with increased neutrophil accumulation in the lungs **(Fig. 6E).** Consistent with the earlier studies [45], *PA* infection led to an increase in the level of proinflammatory cytokines such as *Mip2α* and *Tnfα* in the lungs within 12-24 hours, which reversed to basal levels at 120 hours **(Fig 6F**).

**Figure 6:**
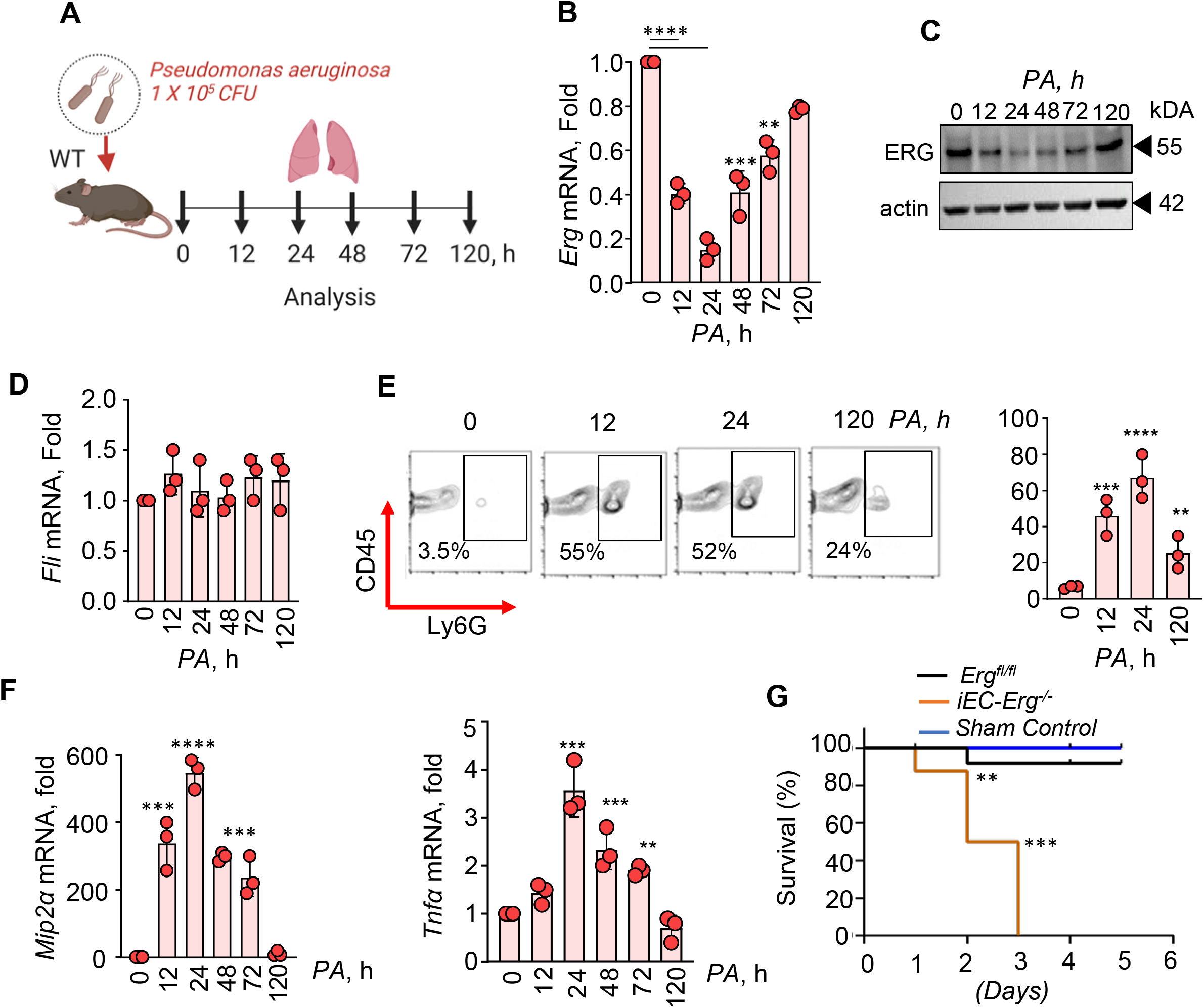
Bacterial pneumonia decreases endothelial ERG and promotes neutrophilic lung injury. **(A)** Schematic of *P. aeruginosa* (PA)-induced lung injury. **(B)** ERG mRNA **(C)** and protein levels at indicated times after *PA* challenge. GAPDH was used as a control for mRNA, while actin was used as a loading control for protein. **(D)** Fli-1 mRNA levels taking GAPDH as control. **(E)** PMN numbers in the PA-infected lungs at indicated times. The left shows the FACS plot, while the right shows quantitation (n=6). **(F)** mRNA normalized to GAPDH. (**G**) Mice received 1x10^5^ CFU of *PA.* Mouse survival was assessed every 6-12 hours after *PA* instillation. n=7 mice/group. Data represented as ±SEM. One-way ANOVA followed by Tukey’s multiple comparison test (all plots). P*<0.05, **<0.01, ***<0.001, ****<0.0001.

Next, we investigated the role of vascular ERG in regulating mouse survival following administration of 1 x 10^5^ CFU of *PA i.t.* in control and *iEC-Erg^-/-^*mice. We assessed the survival of these mice for five days. *PA* infection led to 100% mortality within 72 hours in EC-ERG null mice, while 90% of control mice remained viable for up to 5 days (**Figure 6G**).

## Discussion

Blood leukocytes continuously surveil vascular endothelium to ensure host defense and proper organ function [48]. This function is essential in the lungs, which are continually exposed to environmental pathogens with each breath. The ETS transcription factor, ERG, is well-known for its role in maintaining anti-inflammatory vascular niches [49] [23]. However, how alterations in vascular endothelial function induced by loss of EC-ERG impact PMN function remains unclear. Here, we have delineated the molecular principles governing ERG-dependent PMN function at homeostasis in both murine and human models.

We discovered that PMN infiltration into the airspace at a steady state correlated with a decrease in ERG expression in the EC of mouse lungs. Moreover, a substantial loss of *Erg* during *Pseudomonas aeruginosa* infection in mice and in lungs from ARDS patients was also associated with neutrophil infiltration. PMNs from EC-ERG null lungs were hyperactive, characteristic of inflammatory injury [50]. ERG regulates the expression of several EC genes, including VE-cadherin, and hence plays a crucial role in maintaining vascular homeostasis during development and adulthood [19, 24, 26, 51, 52]. Our current study demonstrates that the number of PMNs in the airspace is tightly regulated and closely linked with ERG’s role in maintaining homeostatic EC.

Our data show that vascular ERG is a unique molecular mechanism regulating the neutrophil transcriptome. PMNs isolated from EC-ERG null lungs showed upregulation of genes linked with pathways inducing inflammation, chemotaxis, and cell survival. We also found increased expression of genes comprising the CXCR4 and CXCR2 pathways in EC-ERG-null PMNs recruited to the airspace compared to control PMNs. CXCR2 in PMNs regulates their chemotaxis [53], while CXCR4 is linked to PMN prolonged survival, pulmonary retention, and disease severity [13]. Crucially, these genes were also expressed in PMNs from ARDS patients. Importantly, we showed that adoptively transferred ERG-null PMNs induced inflammatory lung injury, providing insights into the obligatory role of ERG in suppressing PMN’s transcriptome towards an inflammatory phenotypic switch in the tissue. Loss of EC-ERG leads to vascular hyperpermeability [23, 25], which could induce PMN infiltration and activation [54]. However, we demonstrated that paracrine media obtained from EC-ERG null ECs similarly induced proinflammatory gene signatures in human PMNs.

We identified IL8-CXCR2 signaling as an obligatory mechanism for inducing PMN transendothelial migration and activation in ERG-null lungs. ERG binds to the promoter of several NFkB targets to maintain vascular homeostasis [38, 55]. Inhibition of CXCR2 by Reparixin, an allosteric inhibitor of IL8-CXCR2 chemokine receptor signaling [44], reduced PMNs number in the lungs and subverted PMNs towards the anti-inflammatory lineage, resulting much-improved lung fluid homeostasis in EC-ERG null mice. Reparixin is currently employed to ameliorate neutrophil activation in pulmonary diseases, including bronchiectasis, virus-associated lung infection, chronic obstructive pulmonary disease (COPD), and asthma. Our findings suggest that Reparixin could be a target for preventing vascular inflammation and injury in ERG null or low settings.

Lastly, we demonstrate that A20, a deubiquitinase that regulates multiple functions of ECs [44, 55, 56], underpins the PMN inflammatory phenotype by inhibiting IL8 generation. Data showed reduced A20 expression in ERG null ECs, which was sufficient to augment EC activation and switch PMNs to an inflammatory phenotype. We also showed that ERG binds to A20 promoter and rescuing the deubiquitinase expression and activity in ERG null ECs restored EC lung vascular homeostasis and anti-inflammatory phenotype, reinforcing that the ERG-induced A20 synthesis was the missing arm in ERG-dependent control of EC homeostasis.

In conclusion, our studies show that EC-ERG is an essential suppressor of neutrophilic inflammation (**Figure 7**). As the loss of ERG is observed in chronic lung injury, including fibrosis [23, 57], our findings will have implications for understanding and therapeutically mitigating PMN function in inflammatory diseases.

**Figure 7:**
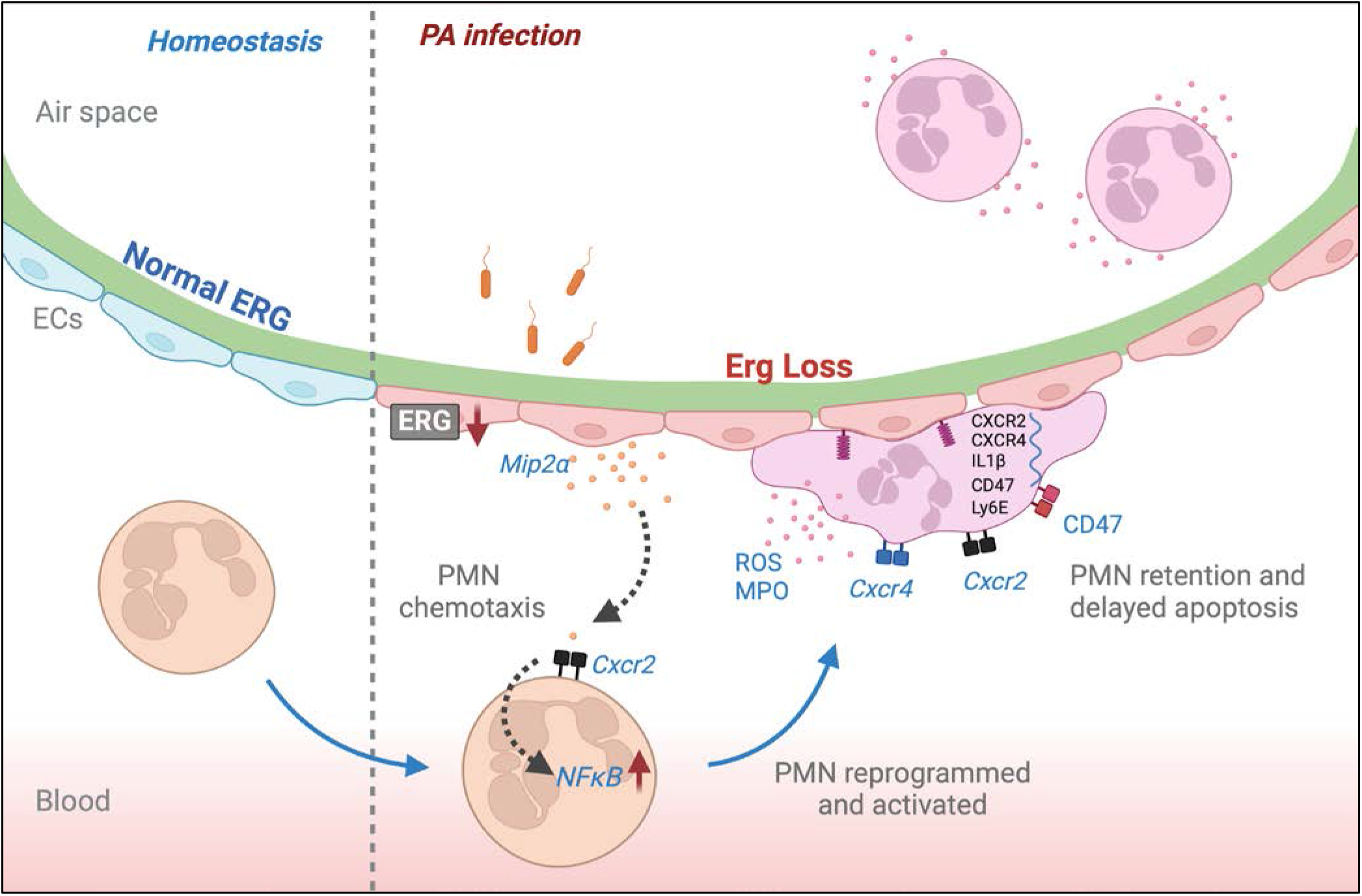
Model of endothelial ERG regulating PMN transcriptome. In normal homeostasis endothelial ERG restricts PMN extravasation and activation by downregulating the NFκB-IL8 cascade. Whereas substantial loss of endothelial ERG during *Pseudomonas aeruginosa* infection rewires neutrophil transcriptome which is associated with inflammation and retention in the lung air space.

### Limitations of the study

Although our study provides compelling evidence for EC-ERG as a crucial mechanism in maintaining the PMN transcriptome, trans-endothelial migration, and activation, we acknowledge several caveats. We show that the genetic loss of vascular ERG phenocopies PMN infiltration seen during pneumonia and in the lungs of ARDS patients. However, future studies are needed to assess the mechanisms that decrease ERG expression in ECs following infections and the transcription factors rewiring PMN identity and activity.

PMNs also undergo reverse transmigration from the tissue to the bone marrow to control tissue homeostasis. While we showed that Reparixin and A20 reduced the number of PMNs in the lungs, it remains unclear whether Reparixin and A20 affect reverse transmigration and apoptosisof PMNs. Furthermore, the details of the mechanisms inducing PMNs to upregulate CXCR2 upon sensing proinflammatory cytokines such as IL8 from ECs are poorly understood.

PMN transcriptomics vary in between homeostasis and inflammation. Understanding the divergent subsets of neutrophils that infiltrate upon ERG loss in ECs remains to be determined. Therefore, exploring ERG-dependent PMNs subsets and defining their tissue-driven transcriptional plasticity during ERG-mediated pulmonary vascular dysfunction may offer insights into PMN-targeted immunomodulatory strategies to prevent various lung complications.

## STAR * METHODS RESOURCE TABLE EXPERIMENTAL MODELS AND SUBJECT DETAILS

### Animals

The Institutional Animal Care and Use Committee of the University of Illinois approved all animal models and experiments used in this study. Tamoxifen-inducible endothelial-specific ERG mice were generated by crossing *Erg^fl/fl^* with mice containing tamoxifen-inducible Cdh5^cre-ERT2^ promoter driver. *Erg^fl/fl^* was a kind gift from Dr Masahiro Iwamoto, The Maxwell Hurston Professor in Orthopaedics, University of Maryland School of Medicine. *Erg*^fl^*/Cdh5^Cre-ERT2^* mice were further crossed with *ROSA-tdTomato* (*Rosa-tomato*: *B6.Cg-Gt(ROSA)*^26Sortm9(CAG-81tdTomato)Hze^*/J*) for converting EC into “tdTomato (red color) [56, 57]. Dr. Yoshi Tsukazaki provided catchup (C57BL/6-Ly6g(tm2621(Cre-tdTomato) mice. LyzM-GFP mice initially provided by Dr. Klaus Ley (La Jolla Institute for Immunology, UCSD) are maintained at the UIC animal facility. C57Blk/6J mice breeding pairs initially obtained from Jackson Laboratory (Farmington, CT, USA) were bred and maintained at the UIC. Mouse colonies were kept in a pathogen-free housing facility at the University. All experiments were performed on C57BL/6J background mice in either gender between 5-8 weeks old.

### Human blood samples

The University of Illinois Chicago’s Institutional Review Board (IRB) approved human blood sample collection and experiments. Briefly, peripheral blood was collected by venipuncture from (n=3) random healthy volunteers and (n=4) ARDS patients. Donors or their surrogates provided written informed consent. Blood PMNs were isolated using MACSxpress® Whole Blood Neutrophil Isolation Kit, human (Cat# 130-104-434). Human lungs from de-identified autopsied patients dying of ARDS or non-lung related diseases were obtained from UIC biorepository. The sections were then immunostained for anti-ERG, anti-Myeloperoxidase, anti-Ly6G antibody and DAPI, followed by immunofluorescent imaging using Zeiss LSM 880 confocal laser scanning microscope as described earlier [58]

### Cell Culture

Human pulmonary endothelial cells (HPAECs) were cultured in a 0.1% gelatin-coated flask using EBM^TM^-2 Endothelial Cell Growth Basal Medium-2 supplemented with growth factors (Lonza # 00190860) and 15% Fetal Bovine Serum (FBS) (Thermo Fisher, # A5256701)[56, 57].

HL60 cells (Human promyelocytic leukemia cells) were grown in suspension with RPMI-1640 medium containing L-glutamine and 25 mM HEPES (Thermo Fisher #72400047), supplemented with 15% FBS (Thermo Fisher #A4766801) and penicillin-streptomycin-amphotericin B (Thermo Fisher #15240096), at 37°C, 5% CO^2^. These cells were exposed to DMSO (1.3%) in RPMI complete media for differentiation and cultured for six days. Differentiated cells were then used for experiments.

### Bacteria

GFP-tagged *Pseudomonas aeruginosa* (GFP-PA01 strain) was used to induce lung injury in mice. [47, 58]

## METHODS DETAILS qPCR

Total RNA was isolated from whole lungs or sorted neutrophils of indicated mice using TRIzol reagent (Invitrogen Inc, Carlsbad, CA) or RNeasy Mini Kit (Qiagen # 74104) according to the manufacturer’s instructions. RNA was quantified using Biodrop, and reverse transcription reaction was carried out using specific primers described in the resource section as per published protocols [56].

### Western blotting

Tissues or cells were lysed using RIPA buffer (Sigma-Aldrich # R0278), and protein concentration was determined using Bradford Assay. SDS-PAGE resolved samples, transferred to 0.45um nitrocellulose membrane (Bio-Rad #1620115) and incubated overnight with primary antibodies. After incubation with a secondary antibody, the protein was detected by chemiluminescence using PICO PLUS (Thermo Scientific, REF 34580) and captured using BioRad ChemiDoc™ Imaging System (Cat# 12003153)[56].

### FACS Analysis

FACS analysis was performed in the indicated lungs as described [56]. Briefly, after mincing, enzymatically digested with 1 mg/mL collagenase A (Roche, New York, NY) for 50 min at 37°C the digested lung tissue was forced through a metal cannula and passed through a 75-mm nylon filter to obtain single-cell suspensions. The red blood cells were lysed using lysis buffer, and the cell suspensions were washed with FACS buffer. Cells were re-suspended in FACS buffer, and after Fc block with FcgRIII/ II antibody for 30 minutes, cells were labeled with indicated antibodies as a cocktail (anti-CD31, anti-CD45, anti-CD64, anti-CD11c, anti-CD11b, anti-Gr1 (anti-Ly6G), anti-SiglecF, anti-CD 62P, and anti-CD3), for 30 minutes on ice. Samples were washed and analyzed using CytoFLEX LX Flow Cytometer (Beckman Coulter), and data were processed using FlowJo™ Software v10.10. All antibodies used for flow cytometry were anti-mouse antigens.

### Immunofluorescence and image analysis

Lungs harvested from indicated mice were fixed in 4% paraformaldehyde (PFA) for four h, after which these were transferred into 30% sucrose solution for 24 h and fixed in OCT FITC-IB4 (50 mg/100 PBS) was injected in each mouse through the tail vein two hours before harvesting lungs. The lungs were cryo-sectioned (4 μm thickness) and immunostained using anti-ERG or anti-VE-Cadherin primary antibodies followed by appropriate secondary antibodies and DAPI. For immunostaining ECs, cells were fixed in 2% PFA for 10 min, followed by with or without permeabilization using 0.1% Triton X-100 for 2 mins. Cells were stained with indicated antibodies and appropriately labeled secondary antibodies and DAPI. Images were analyzed using a Zeiss LSM 880 confocal scanning microscope. All image analyses were quantified by measuring fluorescence intensity using ImageJ software [56, 59].

### Lung intravital imaging and neutrophil assessment

Lung intravital imaging was performed as described [60, 61]. Briefly, *td-Tomato-Erg^fl/fl^* and *td-Tomato-iEC-Erg^-/-^* mice were injected with ketamine (10 mg/ml) and xylazine (2.5 mg/ml) at 40-80 mg/kg body weight (for ketamine) and 10-20 mg/kg body weight (for xylazine).

SeTau647(SETA BioMedicals)-labeled CD31 antibody (25 μg/mice) (clone 390; Biolegend) and BV421-labeled Ly6G (Clone 1A8; Biolegend) antibody (10 μg/mice) was retro-orbitally injected right before the surgery and lung intravital imaging to stain intravascular PMNs and lung microvascular structures. A resonance-scanning two-photon microscope (Ultima Multiphoton Microscopes, Bruker) with an Olympus XLUMPlanFL N 20x (NA 1.00) and Immersion oil (Immersol W (2010); Carl Zeiss) were employed to collect multi-color images (Dichroic mirror; 775 extended pass filter (775 LP; Bruker), IR blocking filter; 770 short pass filter (770 SP; Bruker), Emission filter; 460/50 nm for BV421, 595/60 nm for tomato (Bruker) and 708/75 nm for SeTau647 (FF01-708/75-25; Semrock) with 960 nm excitation at video rate. Motion artifacts were stabilized using computer vision algorithms based on image processing of lung microscopic images. ImageJ and Origin (OriginLab) and customized LabVIEW programs (National instruments) were used to quantify PMN dynamics.

### Bronchoalveolar lavage histology and ROS imaging

Bronchoalveolar lavage (BAL) was performed as described previously [47]. Briefly, after sacrificing the mice, a tracheotomy was performed, followed by the collection of BAL using an 18-gauge blunt needle. The BAL fluid was cytospin by centrifuging at 2000 rpm for 10 min, followed by Hematoxylin and Eosin (H&E) staining to quantify neutrophils. The sections were imaged at 20X magnification using a ECHO Brightfield Imaging at 40 X optical magnification.

The BAL cells plated on a 30 mm cover dish for live ROS assessment were treated with the cell-permeant 2’,7’-dichlorodihydrofluorescein diacetate (H2DCFDA) (Thermofisher Thermofisher #D399). After 30 minutes, the cells were washed and immunostained with anti-Ly6G antibody. dye. After 30 minutes, immunofluorescence was determined by confocal imaging. ROS fluorescence intensity was measured using ImageJ software.

### Transfections

HPAECs were transfected with control (siCont) or ERG siRNA (siERG) targeting Erg-1/2/3 (Santa Cruz Biotechnology, Inc # sc-35334,) (Santa Cruz Biotechnology) to deplete ERG. Briefly, 80% of confluent ECs were transfected with indicated siRNA using siRNA Transfection Reagent (Santa Cruz Biotechnology, Inc # sc-29528,)[56]. Control siRNA (ON-TARGETplus Non-targeting Pool; D-001810-10) was used in all experiments. To rescue A20 expression, ECs transfected with siERG for 24 h were retransfected with A20 WT or A20 mutant cDNA or control vector using Fugene HDTransfection Reagent (Promega # E2311). These cells were then used 24 h after cDNA transfection (Chavez et al., 2015). We performed western blotting and qPCR in all experiments to confirm the ERG depletion or cDNA expression.

### Myeloperoxidase (MPO) assay

Myeloperoxidase (MPO) activity in indicated lung lysates obtained after perfusion with PBS, weighed, and frozen at –80 C for a week was determined calorimetrically using a commercial kit (Abcam # ab105136). MPO activity is expressed as units of MPO activity, where one unit was defined as the change in OD at 460 nm per milligram of protein per minute.

For MPO immunostaining, paraffin-embedded lung sections were stained using recombinant anti-MPO antibody (Abcam # ab208670), followed by appropriate secondary antibody and DAPI. Images were analyzed using a Zeiss LSM 880 confocal microscope. MPO expression is quantified by measuring fluorescence intensity using ImageJ software.

### *Pseudomonas aeruginosa*-induced acute lung injury and survival

*PA* from glycerol-preserved stock was streaked in ampicillin-selective (HiMedia Laboratories LLC, United States) Luria–Bertani (LB) agar plates and incubated overnight at 37° C. Single colonies were picked and inoculated overnight (∼17 h) in 250 ml LB broth containing ampilcillin (100 μg/ml) at 37°C. For the standard plate count method, 1 ml of the bacterial culture from the broth was serially diluted with sterile phosphate buffer up to 1 × 10^10^ dilutions. Each dilution plate was spread-plated in ampicillin selective (100 μg/ml) sheep blood agar plates to get the countable bacterial colonies. The 1 × 10^6^ CFU/25mL was calculated using the following formula: CFU/ml = (no. of colonies x dilution factor)/volume of the culture plate [58, 62]

For lung injury, the anesthetized mice receive 1 × 10^4^ CFU *PA* dissolved in 50 μl PBS/mouse through the endotracheal route. After a 15-minute recovery period, mice were returned to their respective cages. Mice were sacrificed, and the lungs were excised for protein, RNA, FACS, and edema assessment at the indicated times. Edema was measured by determining the lung wet-dry ratio. The left lobe of the lungs was excised following *PA* instillation, weighed, and then dried at 55°C. The dry weight was taken after 24 h, and the wet-dry ratio was calculated, which measured edema, as described previously (Yazbeck et al., 2017).

The acute lung injury was calculated using the method previously described. Briefly, histology of the lung was used to score for (A) PMNs in the alveolar space, (B) PMNs in the interstitial space, (C) hyaline membranes, (D) proteinaceous debris filling the airspace, and (E) alveolar septal thickening. Each item (A-E) was given a score between 0-4. A minimum score of zero would represent a healthy lung, and a maximum score of 1 would represent a lung with severe acute lung injury.

PA (1 × 10^5^ CFU *i.t.*) dissolved in 50 μl PBS/mouse was instilled into mice, and mice survival was assessed every 6-12 hours. The animals’ weight and death were evaluated to determine the survival curve using the Kaplan-Meier graph.

### Neutrophil RNA sequencing

PMNs were sorted using anti-Ly6G, anti-CD45, and anti-CD31 antibodies following labeling of lung cell suspension pooled from three *Erg^fl/fl^* and *iEC-Erg^-/-^* mice. The RNA quality and quantity in sorted cells were assessed using the 2100 Agilent Bioanalyzer. Samples showing RNA integrity number > 8 were only used. Strand-specific RNA-SEQ libraries were prepared using a TruSEQ mRNA RNA-SEQ library protocol (Illumina). Library quality and quantity were assessed using the Agilent bio-analyzer, and libraries were sequenced using an Illumina NovaSEQ6000 (Illumina-provided reagents and protocols). The core analysis was ran in Ingenuity Pathway Analysis on differentially expressed genes to obtain canonical pathways and upstream regulators associated with differentially expressed genes. Significantly enriched pathways were determined based on FDR-adjusted p-value < 0.05.

### Liposome-mediated A20 cDNA delivery to the mouse lungs

Cationic liposomes were made using a mixture of chloroform, cholesterol, and dimethyl dioctadecyl ammonium bromide, as described previously [63]. The liposomes were filtered through a 0.45-micron filter, and A20 cDNA (A20 WT and A20 DUB Mutant) was gently added and mixed. The cDNA-loaded liposomes were administered to control and ERG-deleted mice through the retro-orbital route. After 48 h, the lungs were excised and analyzed for edema, protein, gene expression, and FACS analysis.

### PMN adoptive transfer

PMNs from the lungs of *iEC-Erg^-/-^* and *Erg^fl/fl^* mice were sorted by labeling the lung cell suspension with anti-CD45, anti-CD31 and anti-Ly6G antibodies and 1x10^6^ cells were adoptively transferred into WT or LyzM-GFP mice *i.v*. After 4 hours, the recipient animals were sacrificed, and lungs were harvested to assess the injury and FACS analysis. PBS was used as a vehicle control [47].

### PMN phagocytosis, proliferation, and death

Phagocytic efficiency of PMNs were assessed *in-vivo* by delivering pHrodo™ Red E. coli BioParticles™ conjugate in the *iEC-Erg^-/-^* and *Erg^fl/fl^* mice as per the manufacturers protocol (Thermofisher # P35361). Following this, lungs were excised, immunostained with anti-CD45 and anti-Ly6G antibody and analyzed by FACS. To analyze the PMN proliferation and death, lung suspension from *iEC-Erg^-/-^* and *Erg^fl/fl^* mice were stained with anti-Annexin V and anti-Ki67 antibodies and after 30 min FACS was performed using CytoFLEX LX Flow Cytometer (Beckman Coulter), as described previously [58].

### EC-HL60 co-culture

Differentiated HL60 cells were co-cultured with ECs transfected with control or ERG siRNA. After 24 h, the HL-60 in suspensions were aspirated and collected in a separate tube. The remaining EC adhered PMNs were isolated after trypsinization and using a MACSxpress® Whole Blood Neutrophil Isolation Kit (Abcam Cat# 130-104-434) according to the manufacturer’s protocols. The HL60 cells were then subjected to RNA and protein studies as indicated. HL60 cells adhered to control or siERG ECs were imaged using ECHO Brightfield Imaging at 40 X optical magnification.

### Luciferase reporter assay

Luciferase measurements were performed using the Dual-Luciferase® Reporter Assay System (Promega, # E1910) as per the manufacturer’s protocol. HPAECs were transfected with either empty vector or human A20 wild-type or mutated promoter luciferase reporter followed by stimulation with LPS (1 μg/ml). The Gaussia luciferase (GLuc) and secreted alkaline phosphatase (SEAP) activity were determined as described previously [58]. GLuc activity was normalized taking SEAP activity as an internal control. Luminescence was recorded using a GloMax® 20/20 Luminometer (Cat# E5311, Promega) [56]

### Chromatin Immunoprecipitation (ChIP) Assay

ECs transfected with scrambled or ERG siRNA were stimulated with LPS. Formaldehyde cross-linked protein-DNA complexes (100–125 mg) were immunoprecipitated with the anti-ERG antibody or IgG. DNA fragments were purified using phenol-chloroform-isoamyl alcohol extraction, followed by ethanol precipitation, then resuspended in 15-20 μL nuclease water as a starting material in PCR amplification. The promoter region of A20, which corresponds to a 140 bp fragment containing ERG binding sites, was quantified by real-time q-PCR. Normal rabbit IgG was used as negative antibody control, and DNA from the input (20–40 mg protein-DNA complexes) was used as an internal control [56, 63].

### Statistical Analysis

Results are expressed as means ± SEM from at least three independent experiments. Statistical significance was assessed by one-way ANOVA followed by Tukey’s multiple comparisons test and unpaired t-test with Welch’s correction using Graph Pad Prism version 7.0 (Graph Pad Software, La Jolla, CA).

## Acknowledgments

We acknowledge the UIC-RRC FACS and Histology Cores for their technical support. This work was supported by NIH grants PO1-HL151327, PO1-HL160469, RO1-HL137169, RO1-155941, RO1-HL084153, and RO1-HL165263. We thank Drs Masahiro Iwamoto (Department of Orthopedics, University of Maryland School of Medicine) for the *ERG^fl/fl^* mice, Chinnaswamy Tiruppathi (Department of Pharmacology and Regenerative Medicine, UIC), and Sekhar Reddy (Department of Pediatrics, UIC) for providing the A20 constructs and NFκB signaling antibodies.

## Author contributions

VV and DM designed the study, analyzed the data, prepared figures, and wrote the manuscript. YT performed intravital lung imaging. VV, VBR, and JJ performed mouse experiments. VV, VBR, and SB performed immunohistochemistry of lung tissues and FACS analysis of PMNs. MZA and MA performed an in-silico analysis of transcription factor binding motifs, luciferase reporter assay, and CHIP assay. SD obtained human samples and edited the manuscript; VBR performed RNA-seq data analysis. SP provided key scientific inputs and edited the manuscript; DM acquired funding. All authors edited and approved the manuscript.

## Inclusion and diversity

We support inclusive, diverse, and equitable research conduct.

