## Supplemental-Files for "Endothelial ERG programs neutrophil transcriptome for sustained anti-inflammatory vascular niche"

**Supplementary Figure 1**

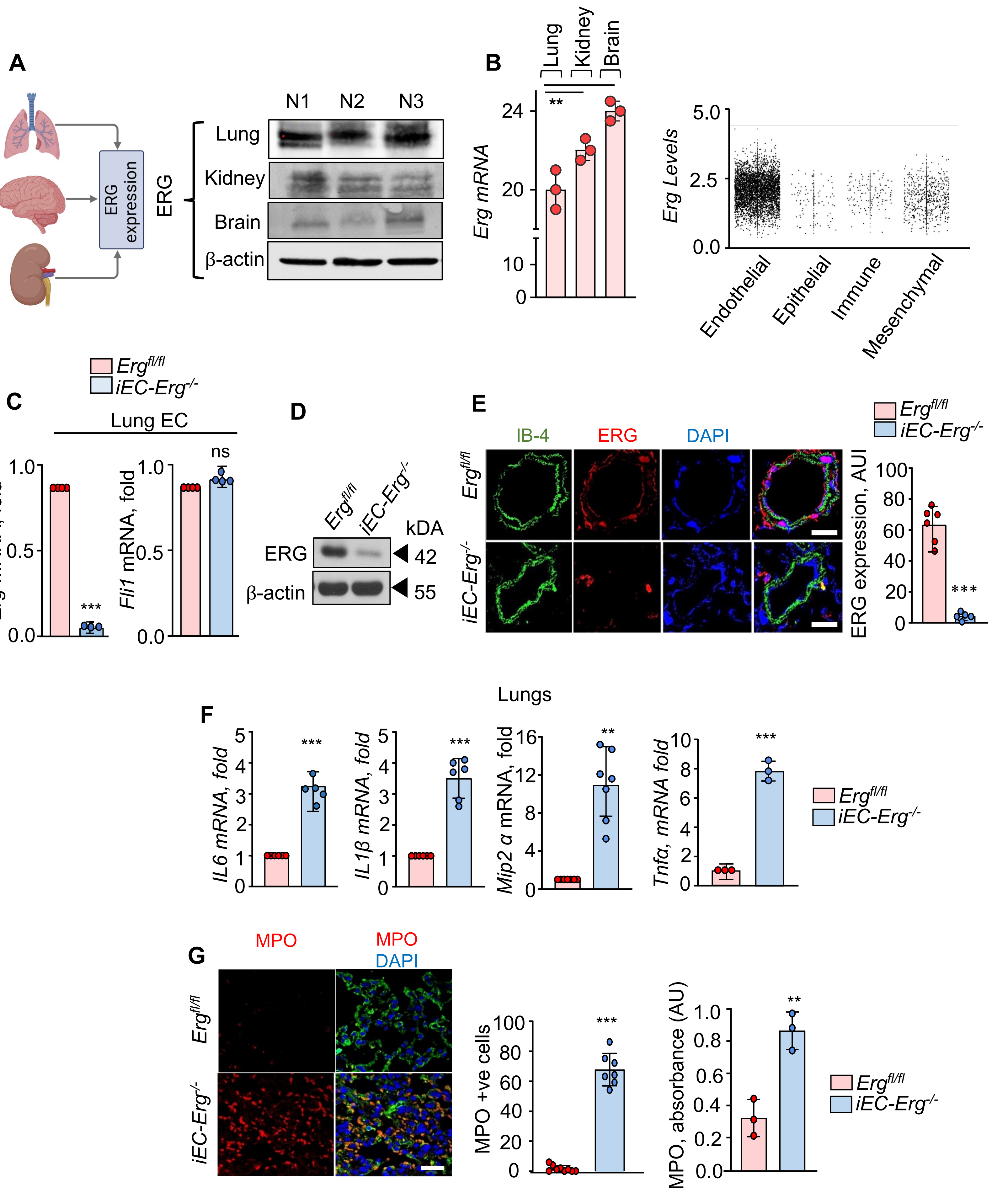

### Supplementary Figure 2

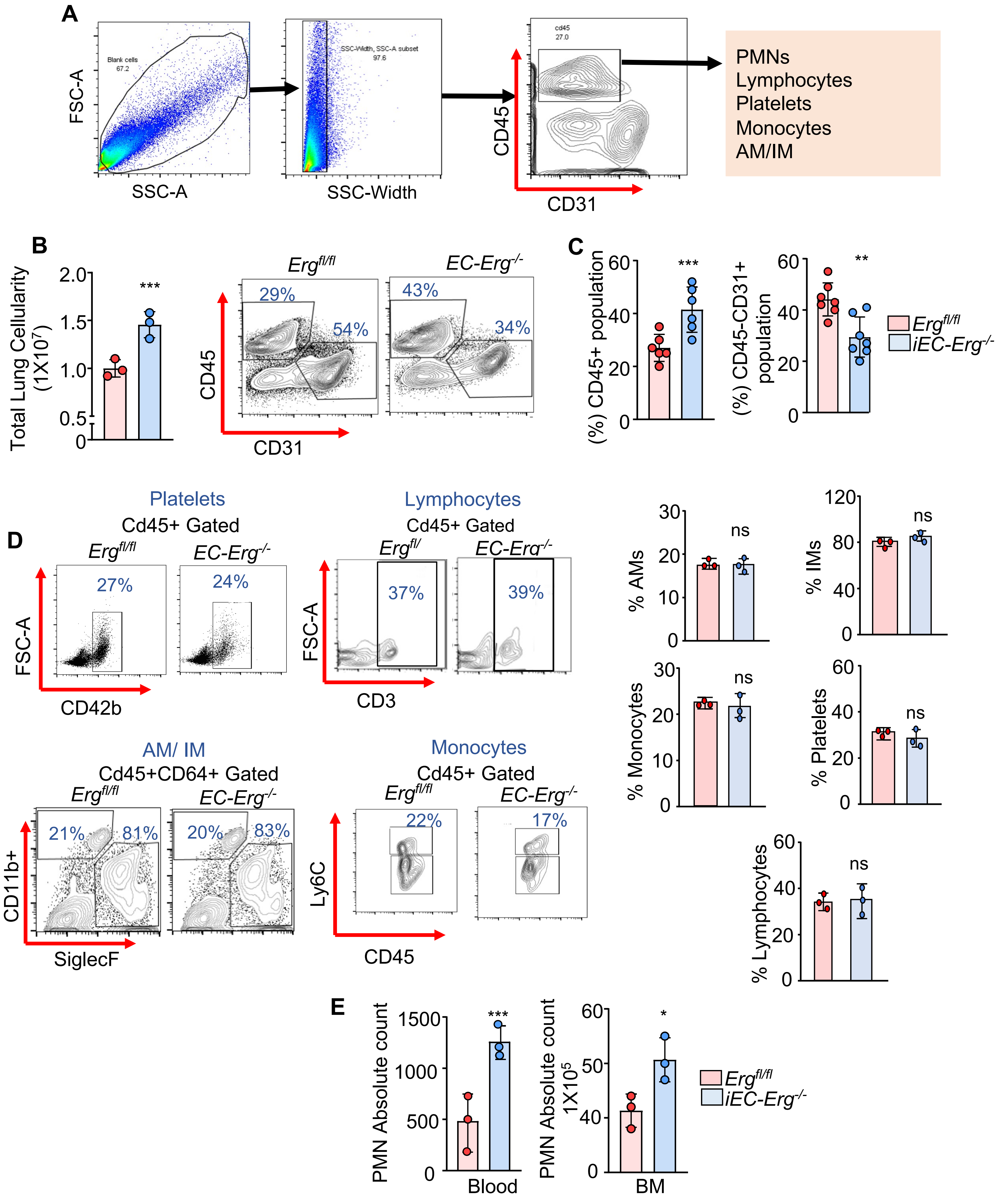

### Supplementary Figure 3

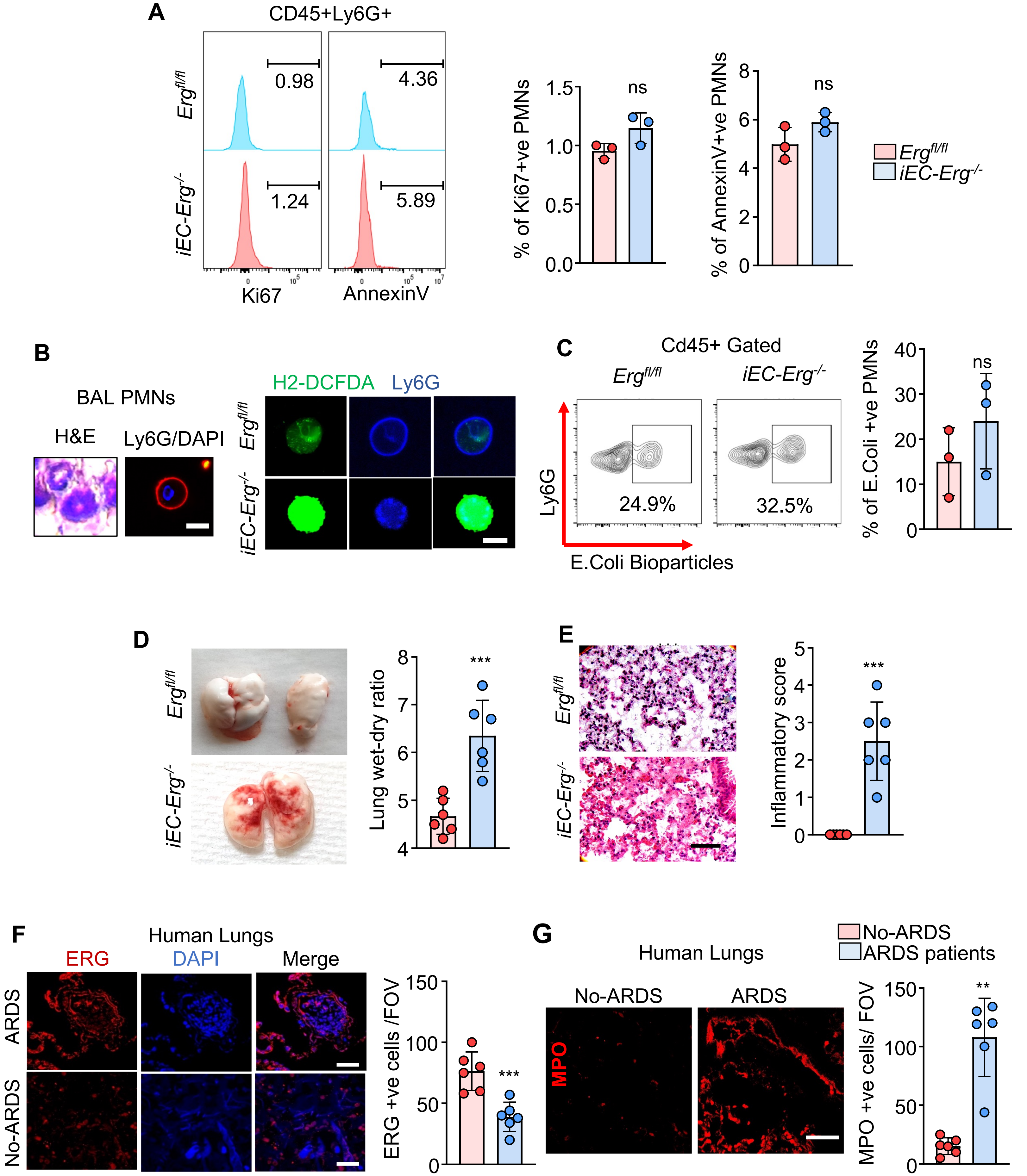

### Supplementary Figure 4

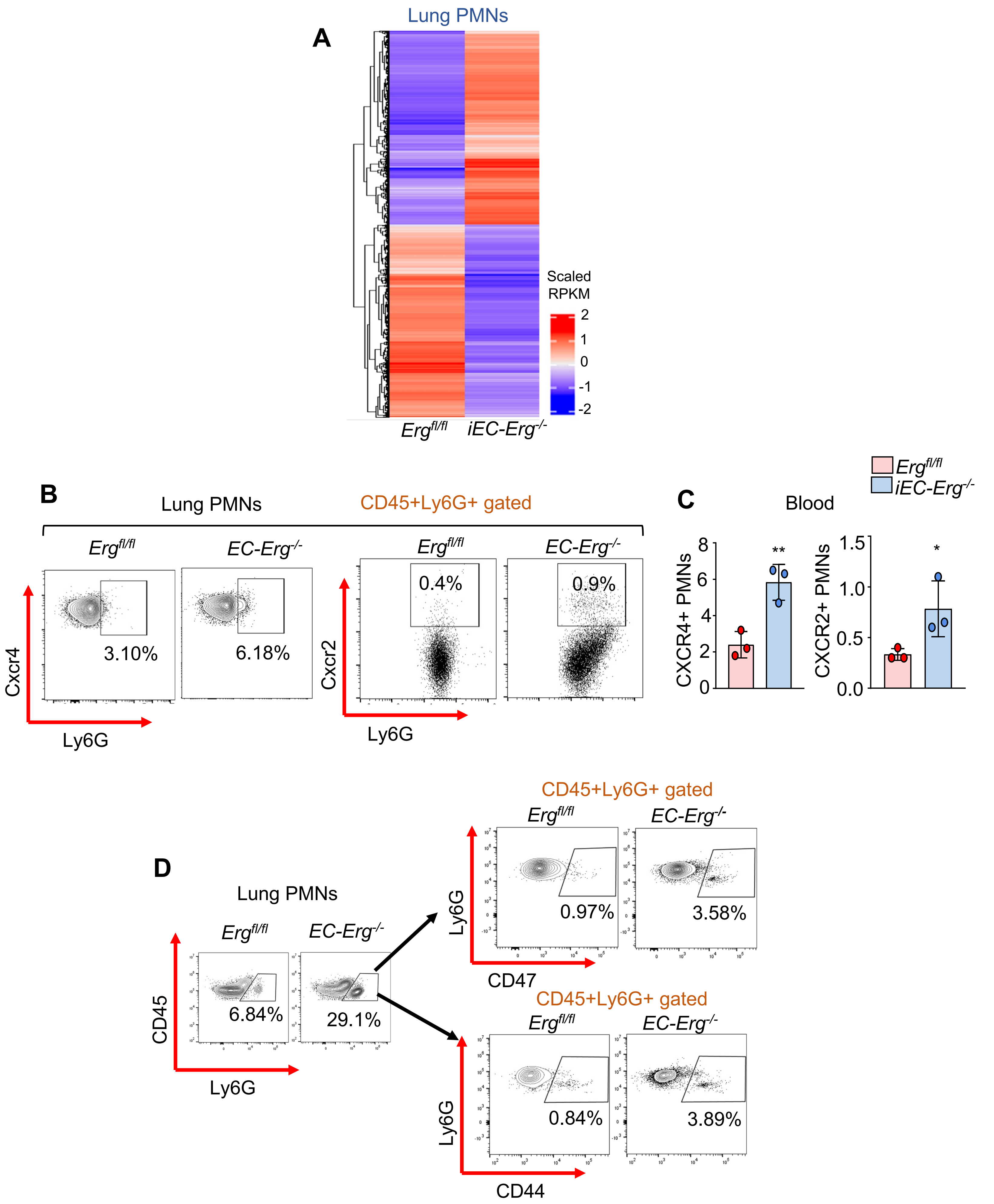

### Supplementary Figure 5

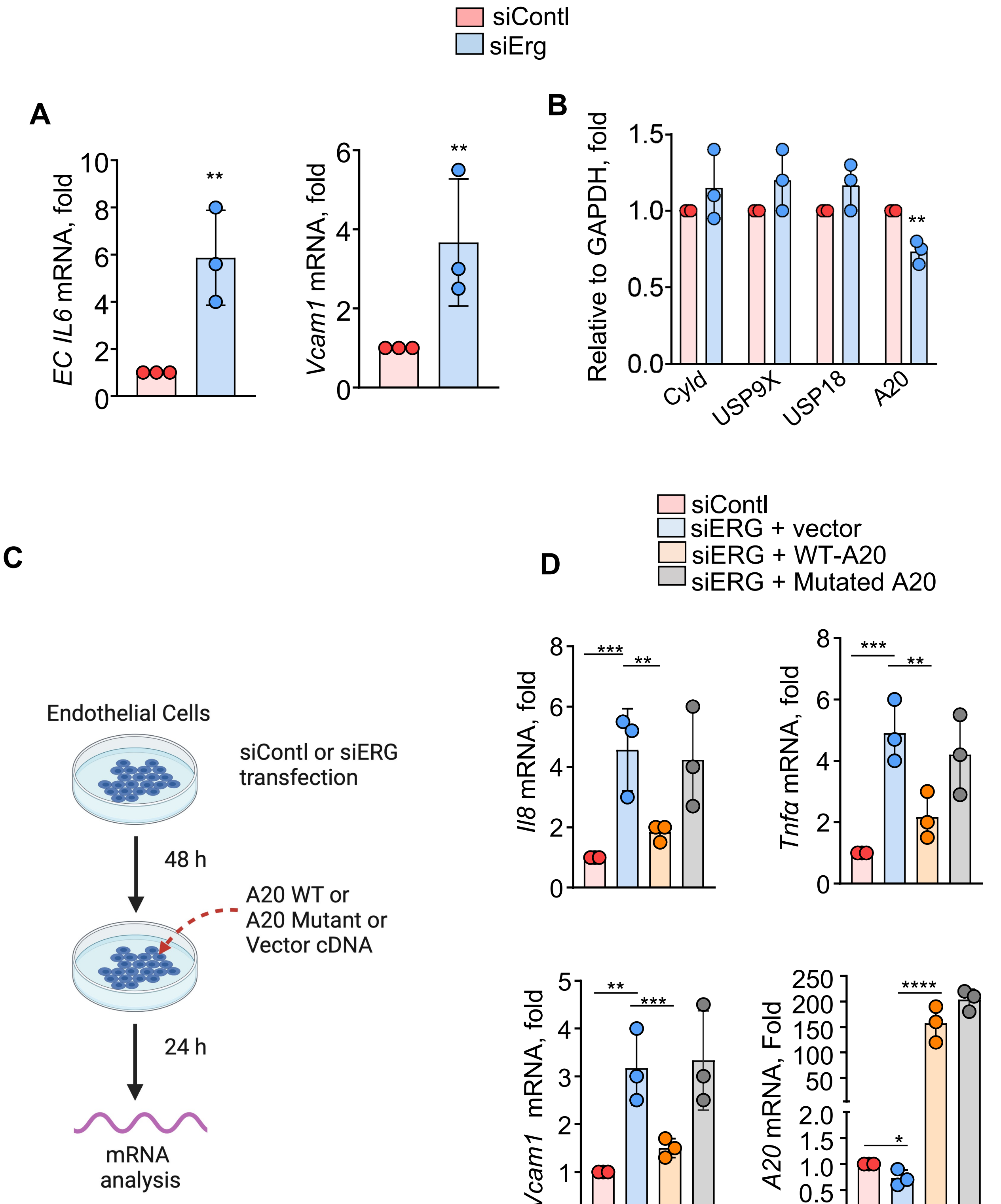

### Supplementary Figure 6

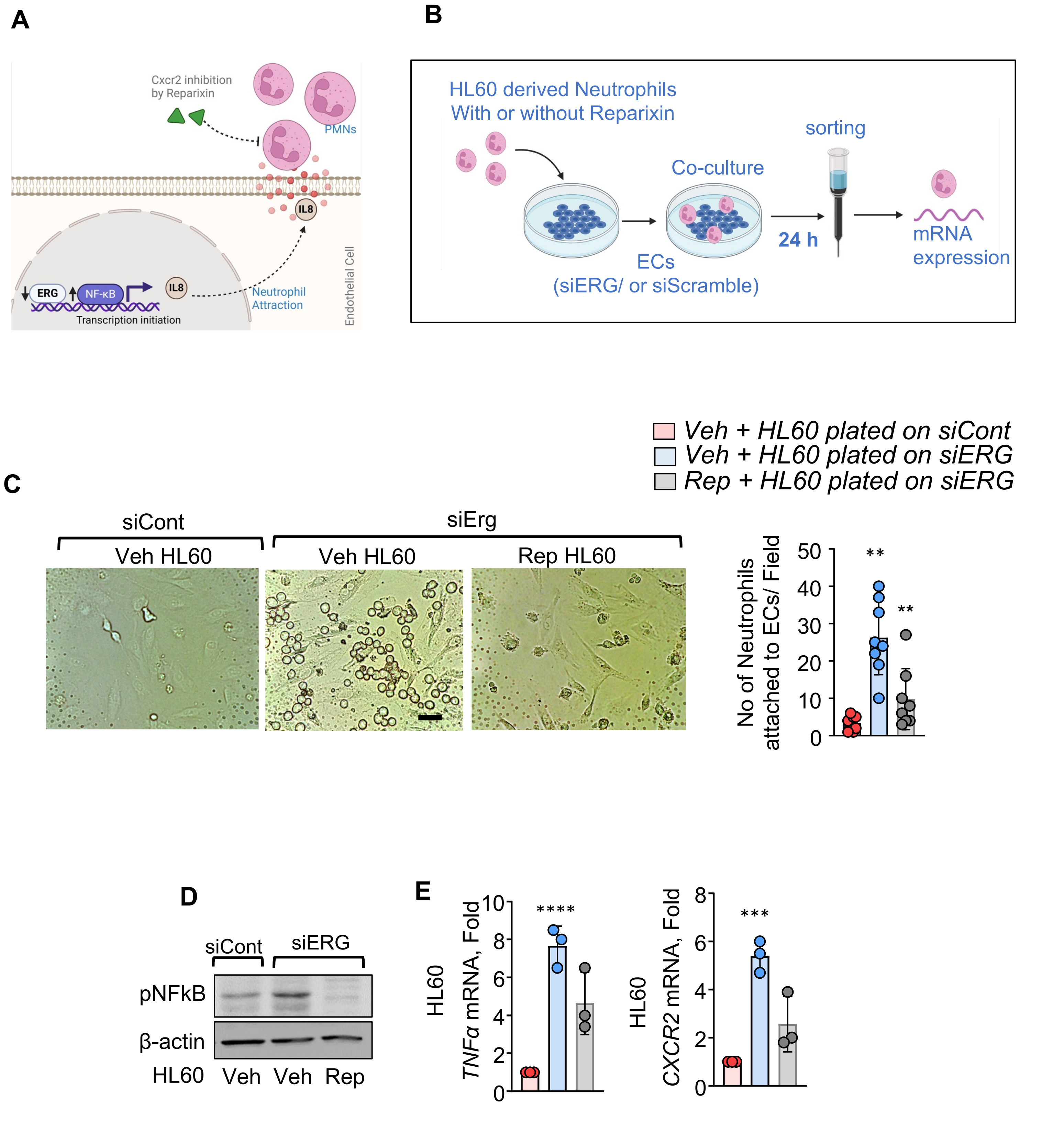
